# Exploring phenotypic and genetic variability in hemp (*Cannabis sativa*)

**DOI:** 10.1101/2023.11.01.565084

**Authors:** Nina Trubanová, Grace Pender, Paul F. McCabe, Rainer Melzer, Susanne Schilling

## Abstract

Hemp (*Cannabis sativa* L.) is a high-value multi-purpose crop with applications ranging from the medicinal use of its secondary metabolites to carbon-sequestering building materials. To fully capitalise on the potential of hemp as a crop for more sustainable agriculture, detailed phenotypic and genotypic characterizations are required in order to inform targeted breeding programmes.

Here, we present a detailed morphological and genomic analysis of 10 hemp cultivars. We found high variability in agronomically important traits such as flowering time, plant height, and biomass in most of the hemp cultivars tested. Additionally, genotyping by sequencing demonstrates that genetic differences are pervasive, both between hemp cultivars as well as between individuals of a single cultivar. The significant genetic and phenotypic variability we observe in hemp contrasts with other crops, where cultivars are often phenotypically and genetically relatively uniform. We argue that the variability of hemp is an asset for breeding and increases the potential for further improvement of the crop but is also a challenge for today’s highly automated agriculture that relies on phenotypic uniformity.

**Author summary:** Hemp (*Cannabis sativa* L.) stands as one of the earliest domesticated crops. This remarkable plant is a sustainable crop with high carbon sequestration capacity which can be cultivated for soil remediation. Furthermore, hemp oil and fibre are used for many applications ranging from cooking to manufacturing bioplastics, textiles, or building materials of superb characteristics, and its secondary metabolites are sought after because of their medicinal properties. However, in contrast to many modern crops, hemp exhibits extensive variability in key agricultural traits, such as plant height and flowering time. This variability presents a challenge for both farmers and processors. To unravel the fundamentals of hemp diversity we conducted a comprehensive study of phenotypic and genetic characterisation of ten diverse hemp cultivars. We present findings confirming substantial variability not only among individuals of different cultivars but also within the same cultivar. Additionally, we explore heterozygosity in the context of other hemp studies and other crops. Understanding this variability in the context of a single hemp cultivar and across multiple cultivars is paramount for breeding novel, more uniform hemp varieties which will allow us to unlock the full potential of hemp as a crop of the future.

## Introduction

Hemp is possibly one of the oldest crops utilised by humans and was likely initially domesticated in the early Neolithic period in East Asia, in what is modern-day China (1). Its fibre was, and still is, used to make textiles, ropes, fishnets, and paper. Meanwhile, the oil extracted from its seeds has served various purposes, including cooking and industrial applications. Additionally, the plant held significance for medicinal and ritual practices (2). Today, the range of utilisation and importance of hemp as a crop constantly increases; indeed within the last two decades, hemp has gained traction as a biomass and fibre crop, as well as for its medicinal uses (3–7).

Because of hemp’s rapid biomass accumulation with low inputs in both conventional and organic farming (8), it has a high carbon sequestration capacity (3) and can be grown as an environmentally beneficial crop. Hemp is also suitable for phytoremediation (9,10), as a defence against erosion (11), and the waste biomass can be processed into biochar which can be used for soil amendment (12). From hemp biomass, it is also possible to produce bioethanol or biogas, and biodiesel from oil extracted from seeds (3,10,13,14). Hemp fibre is used for manufacturing textiles, paper, bioplastics, and biocomposites, as well as construction materials, such as hemp wool used for insulation, hemp clay bricks, or hempcrete (3,5,7–11,13,15,16). Waste products, such as hurd, are used as animal bedding (6,7,11,15,16). Nevertheless, the highest value hemp products are cannabinoids, terpenes, and flavonoids extracted from the hemp biomass, primarily from the female inflorescence, and also from other parts of the plant. Cannabinoids and other secondary metabolites are predominantly used for medicinal and recreational purposes (3–5,15,16), but can also be used as bio-pesticides (6), or for other applications. Hemp oil extracted from seeds is sought-after not only in the cosmetic industry (7,11,13), but because of the ratio of ω-6 to ω-3 fatty acids and protein content (17,18) also as an addition to human and animal diets (4,7,11,16) either in a form of oil or food supplements (3,5). Additionally, besides hemp protein, hemp sprouts and microgreens rich in macronutrients, micronutrients, polyphenols, and antioxidants (4,19) can become a valuable source of nutrients in the human diet.

Meanwhile, novel applications are still emerging, such as functional textiles (10,15) and surgical devices thanks to the antibacterial properties of hemp (15), batteries from carbon nanosheets made of hemp pulp (5), non-petroleum hemp-based ink for tattooing, and hemp filaments for 3D-printing, or oil absorbent materials (5).

Though the plethora of purposes hemp can be utilised for and the importance of this crop is indisputable, hemp research lags behind other major crops. Because of the legal status of hemp (20) in the previous decades little research has been done to maximise the potential of use of this versatile crop. For the implementation of findings in practical applications, including breeding, more relevant hemp variability data must be collected and analysed. Previous studies have begun to explore population genomics and trait variability in order to get a better understanding of the genetics of hemp and to ultimately unlock the potential of this genetically diverse and variable crop (21,22).

While in nature genetic diversity is common among plant species and the basis of evolutionary change (23), in agriculture crops generally exhibit genetic uniformity in their main agronomic traits and this uniformity is favoured by farmers, processors, and consumers (24). Genetic diversity is the sum of the genetic characteristics of a species. In crops, genetic diversity can confer the ability to survive under diverse environmental conditions and in changing climates, as well as provide a genetic reserve of resistance to various biotic and abiotic stressors. It also harbours a collection of genetic variability that is a prerequisite for breeding novel cultivars with improved morphological and agronomical traits (25). Whereas genetic diversity encompasses the distinction of the whole genotypes within the species, variability refers to one or a few different traits of the organism. Genetic variability is observable as distinct phenotypes resulting from contrasting alleles of a gene in individuals of the species or population. Utilising this variability allows breeders to target specific breeding targets.

A strong association of homozygous allelic variations with key agronomic traits has been found in many important crops, such as rice (26), cotton (27), and tomato (28). Among these crops, tomato (29) and certain rice cultivars, particularly its introgression lines (30), have relatively low levels of heterozygosity, while others, like cotton (31), are highly heterozygous. Additionally, other crops like cassava (32,33), mango (34) and grapevine (35,36) were found to be highly heterozygous. Another crop that was found to have high heterozygosity is hemp (*Cannabis sativa*) (21,22).

Here, we assess the phenotypic and genotypic variability of individual plants from ten different hemp cultivars and landraces. Besides intervarietal variability, we find a considerable amount of intravarietal variability, concerning phenotypic as well as genetic traits of the analysed plants. Our results confirm the potential of hemp genetic variability for breeding but caution that at the present individuals of a specific hemp cultivar can differ greatly in one or multiple traits.

## Results

Hemp has been reported to display large phenotypic variability (21,37,38). To explore this variability, ten different hemp cultivars and landraces that varied in flowering time, sex determination, and primary agricultural purpose were cultivated in the greenhouse (Table 1): five monoecious French fibre cultivars that included early flowering ‘Fedora 17’, medium-late flowering ‘Santhica 27’ and ‘Felina 32’, ‘Futura’, and very late flowering ‘Futura 75’ (8,39,40), two accessions of the dioecious hungarian fibre cultivar ‘Kompolti’ (39) with different average flowering time, dioecious oilseed cultivar ‘Finola’, formerly ‘FIN-314’, a cross of two northern Russian landraces (41) that grows well in the arable regions of subarctic Canada and northern Europe (42), a dioecious landrace ‘Georgien’ from the Caucasus region in Georgia, and a dioecious landrace ‘Korea’ from Korea that is genetically more distinct to European and West Asian cultivars (8) (Fig 1a, S1 Fig).

**Fig 1.**
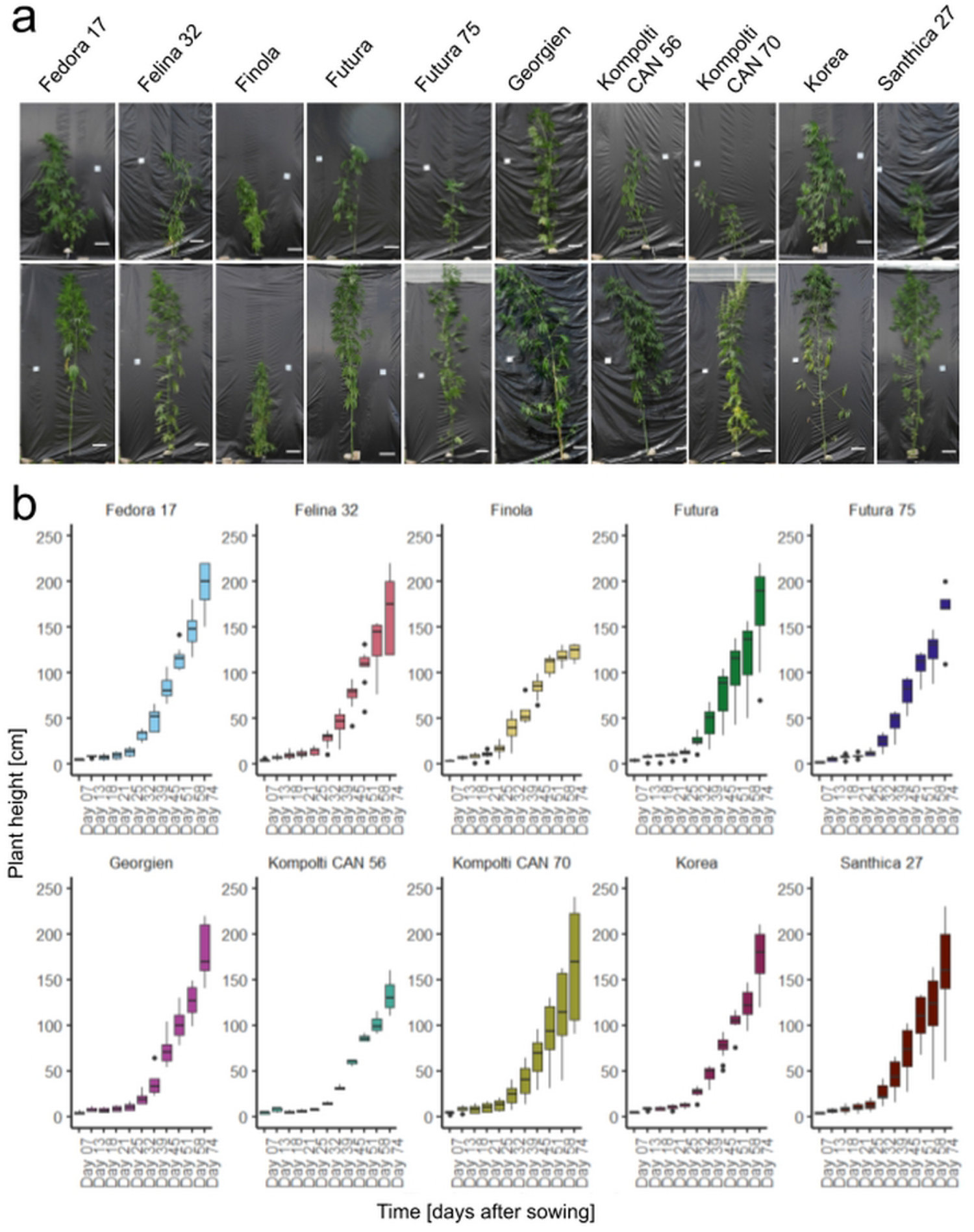
Inter- and intravarietal differences in hemp plant height. The tallest (top) and smallest (bottom) plant per cultivar are depicted (a), scale bars 20 cm. Hemp plant height is represented as box plots measured on days 7, 13, 18, 21, 25, 32, 39, 45, 51, 58, and 74 after sowing. Each panel represents one hemp cultivar (b).

**Table 1.**
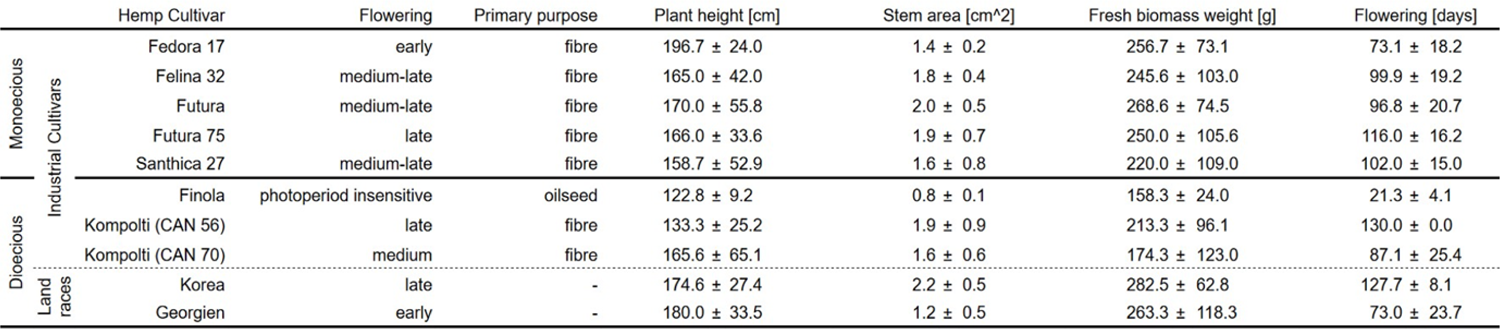
Key traits of ten hemp cultivars grown under greenhouse conditions.

### Hemp cultivars show high variability in biomass and fibre-related traits

Hemp plant height is a good indicator of fibre and biomass yield. Plants grew rapidly over the course of 130 days (Fig 1b). On average, plant height was 163.3 cm over all cultivars. The hemp plants showed high variability in plant height with the tallest individual being four times taller than the shortest individual on day 130 after sowing (Fig 1a). Plant height differences were high, with an average difference between the shortest and the tallest plant within cultivar of 97.2 cm (Fig 2a, S1 Table). On average, the standard deviation of plant height was 36.9 cm, with standard deviations up to 65.1 cm for ‘Kompolti (CAN 70)’, reflecting the variability found in this trait. In some cultivars, such as ‘Santhica 27’ or ‘Futura’, the tallest plant was more than three times taller than the shortest plant observed. Other cultivars, such as ‘Finola’, were more consistent in their height with the smallest individual being 110 cm tall and the tallest being 132 cm tall, and a mean of 122.8 cm (Table 1, S1 Table). Because of large variability within cultivars, significant differences in height between cultivars were only observed between ‘Fedora17’ and ‘Finola’ (Tukey’s HSD, p = 0.039, Fig 2a).

**Fig 2.**
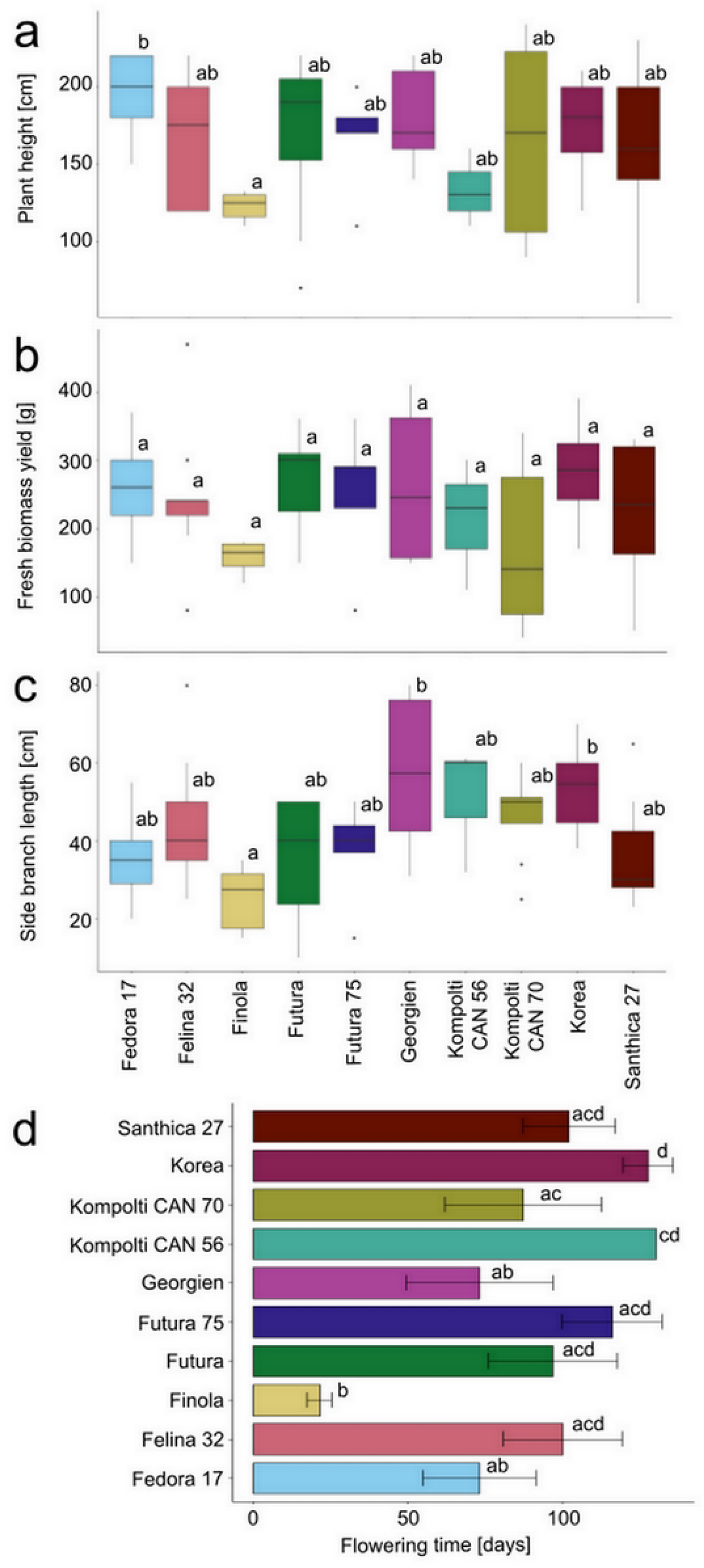
Hemp plant architectural traits and flowering times vary within and between cultivars. Hemp plants were phenotyped for plant height at day 130 after sowing (a), fresh biomass yield (b), and length of the longest side branch (c). Flowering time was observed as the number of days between sowing and the onset of flowering per hemp cultivar (d). Significance levels are indicated with letters (p < 0.05).

Hemp biomass is an important factor for multiple applications, including secondary metabolites extraction (43,44), phytoremediation, and bioethanol production (45), as well as for calculating the carbon sequestration capacity (46). In the hemp plants cultivated, aboveground fresh biomass yield varied almost ten-fold between individuals of the same cultivar, with a mean of 230 g over all cultivars (Fig 2b, Table 1, S1 Table). Large intravarietal and intervarietal differences were observed. No significant differences were observed between cultivars.

Nodes, zones of emerging axillary buds that give rise to side branches, can disrupt the production of hemp fibres, affecting their length negatively (47). The number of nodes per metre determines the quality and length of hemp primary bast fibres and is therefore an important parameter when growing hemp for textile fibre production (47). Between all plants, the number of nodes ranged from four to ten per metre (S2a Fig). Within some cultivars, such as ‘Kompolti (CAN 70)’ or ‘Futura’, the highest number of nodes per metre was twice as high as the lowest number of nodes per metre (S2a Fig, S1 Table) showing the extensive intravarietal variability of this parameter.

Stem area is a good indicator of fibre yield if sowing density is optimised (48–50). The stem area 5 cm above ground ranged from 0.38 to 2.99 cm^2^, with a mean of 1.94 cm^2^ (S2b Fig, Table 1, S1 Table). The mean stem area of the landrace ‘Korea’ of 2.24 cm^2^ was more than twice as large as the stem area of ‘Finola’ of 0.81 cm^2^ (Table 1, S1 Table).

Plant architecture within the *Cannabis* genus is highly variable. To approximate plant architecture, the length of the longest side branch was measured. We observed that some individuals were slender while others were considerably wider (S1 Fig), with the longest side branches ranging from 10 to 80 cm (Fig 2c, S1 Table). The highest intravarietal differences in the length of the longest side branch were measured in ‘Felina 32’ with an average difference of 55 cm between individuals (Fig 2c, S1 Table).

### Hemp cultivars show high variability in flowering-related traits

All studied cultivars, with the exception of ‘Finola’ which is a photoperiod-insensitive cultivar, are short-day plants. Besides day length, hemp flowering time is affected by environmental conditions. Hemp cultivars grown in warmer climates with a higher number of cumulative sunlight hours tend to flower earlier and over a longer period than when grown in colder climates (51). Moreover, male individuals flower and senescence earlier than female individuals (52,53). ‘Finola’ male plants took 19 days or less to flower. In female ‘Finola’ plants, flowering time varied between 24 and 33 days after germination. For all other hemp cultivars, flowering time varied between 50 and 130 days (Fig 2d, Table 1, S1 Table). Late-flowering cultivars, namely ‘Futura 75’ and ‘Kompolti CAN 56’, and ‘Korea’ landrace were the latest flowering hemp cultivars. Some individual plants of the monoecious cultivars ‘Futura 75’ and ‘Santhica 27’ did not flower at all prior to the harvest on day 130, underlining a vast intravarietal variability within flowering time.

Inflorescence density is a good predictor of future seed and potential cannabinoids, terpenes, flavonoids, and sterols yield (54,55). The variability of flower density was higher in monoecious cultivars, such as ‘Felina 32’, ‘Futura’, ‘Futura 75’, and ‘Santhica 27’ (S3 Fig). The highest flower density was observed in female individuals of the cultivar ‘Finola’, which is grown for flowers and seeds.

Hemp is a naturally dioecious plant with male and female flowers located on separate plants (56). Monoecious cultivars with male and female flowers on the same plant that originated from breeding are also available and favoured for some applications (57,58). Large sexual dimorphism affects phenotypic traits and plant morphology. In three out of five monoecious cultivars, ‘Fedora 17’, ‘Felina 32’, and ‘Futura’, only female flowers and no male flowers were observed in single individuals (S4 Fig).

### Individual hemp plants are genetically variable

We detected substantial inter- and intravarietal phenotypic variation for every trait analysed. Therefore, we aimed to understand whether this phenotypic variation is mirrored at the genetic level. We employed genotyping-by-sequencing of 22 selected plants (S2 Table) to investigate single nucleotide variants (SNVs) within and between cultivars.

The number of SNVs per individual varied between 407,799 and 485,430 SNVs (Fig 3a, S2 Table). The number of SNVs per chromosome was significantly different between individuals (Chi-square, p-value < 2.2e-16 for all individuals). Visualisation of the density of SNVs showed that SNVs are overall unevenly distributed on all chromosomes of individual hemp plants (Figs 3b and 3c, S5 Fig). In general, density of SNVs was higher at putative subtelomeric regions and low in putative centromeric regions (59).

**Fig 3.**
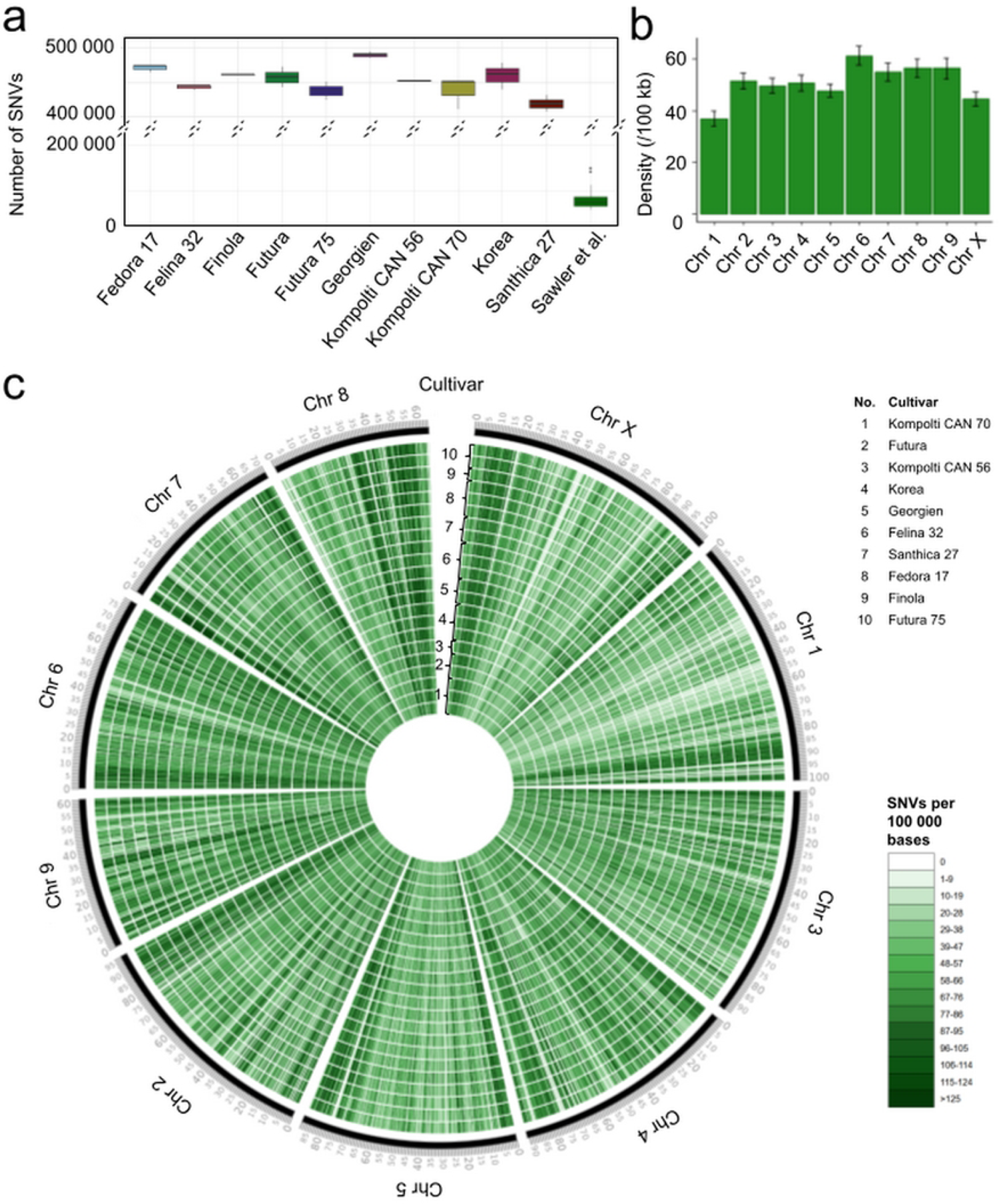
Short nucleotide variants (SNVs) of ten hemp cultivars. The number of filtered SNVs generated by genotyping-by-sequencing per hemp cultivar and of hemp cultivars previously published (21) were visualised as boxplots (a). The mean density of SNVs of all cultivars was visualised as barplots with SD per chromosome (b). The distribution of SNVs for each individual plant was visualised as a circos plot (CircosVCF, 79) (c). The density of SNVs (b) and their distribution (c) are represented per 100,000 bp using the *C. sativa* reference genome (93).

The genetic variability of the studied plants was further examined by calculating their individual heterozygosity based on the ratio of heterozygous to homozygous SNVs (Fig 4a, S3 Table). The heterozygosity on average was 0.269, ranging from 0.189 in one of the ‘Santhica 27’ individuals to 0.338 in one of the ‘Kompolti CAN 70’ individuals. This is similar to previously published data (Fig 4b, S3 Table) (21,22). ‘Santhica 27’ displayed the lowest average heterozygosity (0.216). Conversely, the ‘Futura’ hemp cultivar exhibited the highest average heterozygosity (0.313). To contextualise this finding, we compared the observed heterozygosity in hemp to SNV data of thousands of accessions of 10 other crop species (S6 Fig). Crops with an average heterozygosity higher than collated hemp data (mean: 0.269, median: 0.267) were *Brassica napus* (mean: 0.543, median: 0.490), *Cucurbita pepo* (mean: 0.458, median: 0.439), and *Cucumis sativus* (mean: 0.376, median: 0.286). A number of cultivars from other species had a considerably lower heterozygosity than hemp cultivars, including watermelon (*Citrullus lantanus*), soybean (*Glycine max*), cotton (*Gossypium hirsutum*), and common bean (*Phaseolus vulgaris*). Also, some improved or inbred lines from maize (*Zea mays*) and wheat (*Triticum aestivum*) had very low levels of heterozygosity.

**Fig 4.**
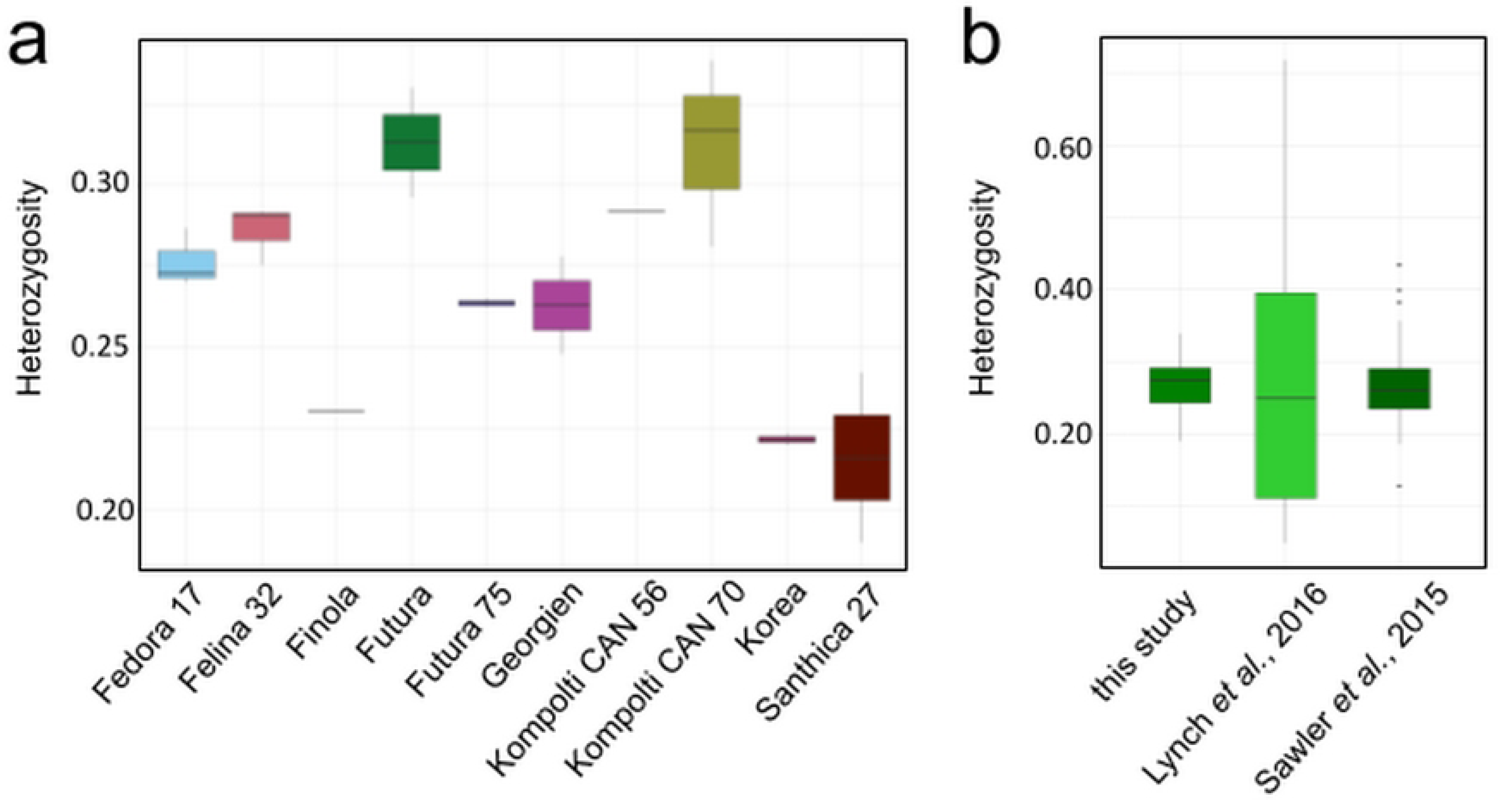
Hemp genomes have a high degree of heterozygosity. Variability in heterozygosity of ten hemp cultivars was calculated as a ratio of heterozygous to homozygous alternative alleles for ten hemp cultivars (a) and other publicly available hemp data (21,22) (b). Each VCF file from Sawler *et al.* (2015) (21) data and Lynch *et al.* (2016) (22) data was intersected with a VCF file generated with FreeBayes (79,80) from our collection merged in the output to a single file using VCF-VCFintersect (83) to use limit analysis to sites present in GBS data. The resulting VCF files contained only sites identical to those found in our data.

### Individuals from the same hemp cultivar differ significantly genetically

Observed phenotypic differences and differences in the number of SNVs and heterozygosity levels between hemp plants of the same cultivar were in many cases as high as between cultivars. Therefore, we analysed whether hemp plants can be associated with a specific cultivar based on the SNV genotyping. A principal component analysis (PCA) analysing SNV profiles of individual hemp plants based on the variance-standardised relationship matrix showed a strong trend of clustering for individuals belonging to the same variety (Fig 5a, S7 Fig). Individuals from the two accessions of ‘Kompolti’ (CAN 56 and CAN 70) created one cluster, similarly individuals of ‘Futura’ and ‘Futura 75’ clustered together. A large cluster was created by plants belonging to monoecious French hemp cultivars ‘Fedora 17’, ‘Felina 32’, and ‘Santhica 27’, and dioecious cultivar ‘Finola’ (Fig 5a). Dendrograms visualising results of different types of clustering, specifically based on pairwise dissimilarity (Fig 5b), pairwise identity-by-state (S8 Fig), and hierarchical hclust clustering (S8 Fig), confirmed results of the PCA and close genetic relationship of hemp plants belonging to the same cultivar.

**Fig 5.**
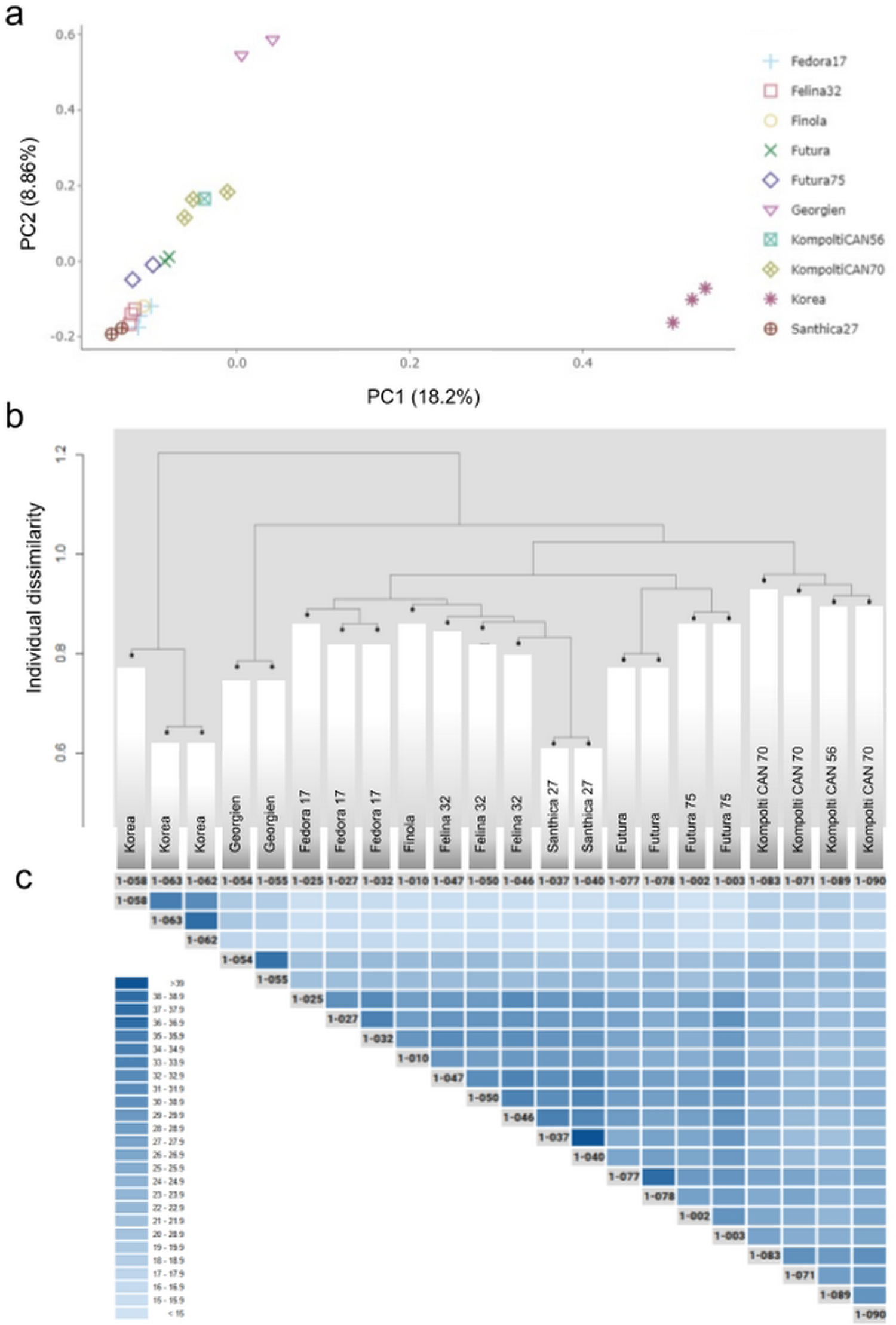
Individual hemp plants from the same cultivar show a high degree of similarity. A principal component analysis (PC1 and PC2) shows a clustering of individuals of the same hemp cultivar (a). For PC1 and PC3, and PC2 and PC3 see S7 Fig.

A hierarchical cluster dendrogram based on pairwise dissimilarity demonstrates clustering of individuals from the same cultivar (b). Relative amounts of alternative SNV positions overlap between individuals show an increased similarity between individuals from the same cultivar (c). The percentage of shared SNVs is expressed as a ratio of shared SNVs to all SNVs of both individuals combined in percentage; i.e. if individuals share 100% of positions the value is 100, if they share 50% of positions the value is 33.3.

The large amount of differences in the number of SNVs and heterozygosity of individuals prompted us to investigate whether individuals differed in their SNV positions. The chromosomal positions of SNVs (hetero- and homozygous) in each sequenced individual were compared to SNV positions of all other individuals. The number of shared SNVs was then expressed as a ratio to all SNVs of the two individuals as percentage (Fig 5c). Individuals of the same cultivar showed an overlap, but not a full congruence of SNV positions (Fig 5c). For example, between two ‘Felina 32’ individuals, only 32.09% of SNV positions were identical. However, the percentage of SNV positions identity between individuals of the same cultivar was overall higher than between individuals of different cultivars. European cultivars had a higher percentage of shared SNVs (Fig 5c) than the landraces ‘Korea’ or ‘Georgien’ which is in line with the results of the PCA and clustering (Fig 5a).

## Discussion

In this study, we conducted a comprehensive phenotypic analysis involving more than 80 individuals representing ten distinct hemp cultivars. To unravel the genetic variability of these cultivars we genotyped 22 of these individuals using genotyping-by-sequencing.

Our data show that individuals within a given cultivar exhibited a higher degree of genetic similarity to one another when compared to individuals from different cultivars. The relationships constructed on the basis of our sequencing data align closely with documented breeding histories, for instance, the close relationships of French fibre cultivars.

### Hemp phenotypic traits are diverse, both between as well as within cultivars

Hemp is a fast-growing crop which accumulates biomass rapidly (Figs 1 and 2a). Therefore, hemp has a huge potential as a carbon-sequestering crop, with a multitude of industrial applications (3). Its short growth cycle makes it particularly interesting as a break crop since it has soil-remediating properties (9,53).

While hemp cultivars are firmly established and used in both agriculture and industry, we found that specific cultivars often do not significantly differ in agronomically important traits, including plant height and biomass yield, plant architecture, and flowering time (Fig 2). In fact, for most traits we investigated, the variability among individuals within the same cultivar was so pronounced that significant differences between cultivars could not be detected (Fig 2, S2 Fig). Consequently, while certain general trends can be observed among hemp cultivars, the extensive intravarietal variability in all the observed phenotypic traits often makes it impossible to classify an individual plant based on our data as belonging to a specific cultivar. It is worth noting that other traits not addressed in this study, such as cannabinoid content or fibre quality, may or may not exhibit similar levels of variability (13,32).

### Hemp genomes display a high degree of heterozygosity

We hypothesise that the substantial phenotypic differences among individuals are, to a significant extent, genetically determined. Our SNV genotyping reveals a relatively high degree of individual heterozygosity. This heterozygosity implies that offspring will exhibit variability in genetically determined traits controlled by genes that are heterozygous in the parental plants. While a certain degree of heterozygosity offers advantages for plant populations (24,61), homogeneity is desirable in agricultural and industrial contexts. In the case of drug-type *C. sativa* (marijuana), the industry standard involves working with clonal propagation of mother plants that exhibit the desired phenotype, rather than relying on seed propagation. This approach is generally impractical for field-grown, high-volume crops like fibre hemp cultivars of *C. sativa*. Reducing the heterozygosity of hemp cultivars through breeding efforts introduces a potential avenue to increase the profitability of the entire value chain by improving harvesting and processing steps.

Conversely, selfed populations characterised by low heterozygosity can suffer from inbreeding depression (62), manifested in reduced yields compared to cross-pollinated progeny (63). Hence, a certain level of heterozygosity may be desirable. However, achieving this balance requires the development of low heterozygosity parental lines and the optimisation of the subsequent seed production in the context of hemp cultivation. *C. sativa* is primarily a dioecious plant, although monoecious populations exist as well (64). In monoecious cultivars, lower hemp heterogeneity resulting from autogamy (65) allows the harvest of both seeds and stems for fibre (54). Self-pollination has proven effective as a method for determining the presence and levels of cannabinoids in autogamous populations. It also serves as a breeding technique to create starting material with stable absence or presence of these compounds (66). Interestingly, in our study, both dioecious and monoecious cultivars exhibited comparable levels of heterozygosity (Fig 4a).

We compared the SNV positions among sequenced individuals. Our findings revealed that similarities were most pronounced among individuals of the same cultivar, followed by individuals of closely related cultivars. The lowest number of SNV positions overlapping were detected between individuals of dissimilar cultivars (Fig 5c). The relationships derived from our sequencing data (Fig 5b) corroborate known breeding histories. For instance, French fibre cultivars such as ‘Felina 32’, ‘Santhica’, and ‘Futura’ (39) are more closely related to each other than to landraces ‘Korea’ and ‘Georgien’. The higher density of SNVs in distal parts of the chromosomes and lower density in centromeric regions (Fig 3c) is in consensus with a high recombination rate at chromosome ends and reduced rates in the putative centromeric regions (67).

High heterozygosity levels are similar to other highly heterozygous crops such as cucumber and *Brassicaceae* (S6 Fig). The heterozygosity in hemp is however higher than that of many crops, including maize and soybean, where even landraces do not possess heterozygosity levels as high as observed in hemp. Numerous studies have previously described high rates of heterozygosity, especially in *Brassicaceae* and *Cucurbitaceae* species, and linked them to diverse phenotypes between cultivars of the respective crop species (68–71). In *Brassicaceae*, the high level of heterozygosity in commercial cultivars stems from the generation of F1 hybrids (71). High heterozygosity as a sign of heterosis effects is one strategy for crop improvement in these crops, while in other cases, high heterozygosity might be counterproductive (72).

### Genetic and phenotypic variability presents both opportunities and challenges in hemp breeding

The extensive range of phenotypic trait values and the underlying genetic heterogeneity makes farming more challenging. To illustrate, variations in flowering times among individuals of the same cultivar (Fig 2d) inevitably impact the timing of harvest and processing for inflorescences and seeds. Therefore, achieving uniformity in traits related to hemp inflorescences, such as flowering time, inflorescence density, and consistent sex expression in monoecious cultivars, should be a central focus of breeding efforts. Furthermore, for research and genetic analyses, sequences from inbred individuals can be utilised to improve the current genomic assemblies (38) that are prone to misassembly due to heterozygosity (73).

We observed significant variability not only among individuals from different cultivars but also among individuals of the same cultivar. Consequently, when conducting comparative studies involving different hemp cultivars, as well as cultivars of other highly heterozygous crops, relying on a single plant (21,22,74) or just a few individuals (16,66) as representative samples is insufficient.

To expand the *C. sativa* gene pool, a comprehensive genomics-based assessment of the *C. sativa* germplasm is essential to define the available diversity (73). This approach would facilitate the identification of superior alleles contributing to hybrid vigour and the development of elite cultivars, much like what has been achieved in rice (4). Our research highlights the substantial diversity within hemp genomes, offering valuable resources for both breeding and research. This genetic wealth can be leveraged to develop novel cultivars that exhibit high-value characteristics and uniformity in traits crucial to various applications. Hemp’s notable phenotypic plasticity positions it favourably for successful cultivation across diverse climates and environmental conditions.

## Materials and methods

### Plant growth

Ten cultivars of hemp (*C. sativa*) were grown in a greenhouse between May and September 2019. Seeds of ‘Futura’ (CAN 69), ‘Georgien’ (CAN 22), ‘Korea’(CAN 23), and both accessions of ‘Kompolti’ (CAN 56 and CAN 70) cultivars were obtained from IPK Gatersleben, and seeds of ‘Fedora 17’, ‘Finola’, ‘Futura 75’, ‘Felina 32’, and ‘Santhica 27’ cultivars were obtained commercially. Plants were grown in 2L pots with a custom soil mix of 1:1:1 John Innes No. 2:Vermiculite:Perlite and fertilised once a week with water-soluble fertiliser (NPK 24-8-16, Water Soluble All Purpose Plant Food, Scotts Miracle-Gro Products, Inc.). The position of plants was randomised and plants were rotated multiple times during the trial to avoid bias associated with their position in the greenhouse.

### Phenotypic measurements and records

Plant height was measured at several time intervals during the plants’ growth on 7 days, 13 days, 18 days, 21 days, 25 days, 32 days, 39 days, 45 days, 51 days, 58 days, 74 days, and at harvest. Plant height was measured from the base of the plant to the top of the main stem. Flowering was defined as the presence of a female flower with visible stigmata or a male flower bulb with a length of at least 3 mm. The day when flowers were first observed was recorded as flowering time for each plant expressed as a number of days after sowing. After day 67, flowering was observed on day 74, day 102 and day 130, hence observed flowering times are upper estimates.

Each plant was sexed and recorded as dioecious if being either male or female, or monoecious if both, male and female flowers, were observed on the same plant. Data related to other phenotypic parameters, such as flower density, number of nodes per one metre, longest side branch length, stem area, and above-ground biomass were collected at the time of the harvest. Flower density was subjectively categorised using the ‘low’, ‘medium-low’, ‘medium’, ‘medium-high’, and ‘high’ density scale. The number of nodes per one metre was taken from the bottom of the stem. Stem surface was calculated using ImageJ from aerial photos of plant stems cut ∼5 cm above the soil line.

### Tissue harvesting and DNA extraction

After about eight weeks, tissue samples were harvested from the young leaves. Samples were snap-frozen in liquid nitrogen and stored at −80°C before DNA extraction. DNA was extracted from each sample using a DNeasy Plant Mini Kit (Qiagen, Germany). DNA was quality-controlled using gel electrophoresis and photospectrometry.

### Sequencing and data acquisition

Sequencing data were obtained using genotyping by sequencing (GBS) method essentially as previously described (74), but DNA was digested with MseI and the sequencing platform used was Illumina HiSeq. Paired-end 150 bp sequencing was performed (CD Genomics, USA). The raw sequence has been deposited in the NIH Sequence Read Archive (SRA), under BioProject PRJNA1023932. Publicly available genotyping by sequencing (GBS) data (21) and whole-genome shotgun sequencing (WGS) data (22) for hemp were obtained from the NCBI. SNV data for other crops were retrieved from the Plant Imputation database (75).

### Mapping and variant calling

A collection of trimmed sequences was aligned to cs10 *C. sativa* reference genome assembly GCF_900626175.2 using the Burrows-Wheeler Alignment (BWA-MEM) tool (76–78). Variants were called with FreeBayes (79,80) on each file individually for calculating heterozygosity and SNV densities, and on all files together for PCA, dendrograms, and SNVs positions comparison. All analyses were performed using the platforms UseGalaxy.org and UseGalaxy.eu (81).

### Heterozygosity analysis

For calculating heterozygosity, variants were filtered for quality > 30 and read depth > 5 or 9 for GBS and WGS respectively using bcftools annotate (82) and VCFfilter (83). Out of all types of called variants, only SNVs were kept using bcftools annotate (82). Homozygous reference sites were removed using the bcftools filter (82). Each VCF file from Sawler *et al.* (2015) (21) data and Lynch *et al.* (2016) (22) data was intersected with a VCF file generated with FreeBayes (79,80) from our collection merged in the output to a single file using VCF-VCFintersect (83) to use limit analysis to sites present in GBS data. The resulting VCF files contained only sites identical to those found in our data. The heterozygosity of each plant was calculated as a ratio between the number of genotyped heterozygous SNVs and the number of non-reference homozygous SNVs using HetHomAlleles (83). The same tool was used to calculate individual heterozygosity from SNV data obtained from the Plant Imputation database (75).

### PCA and clustering

Firstly, variants were filtered for quality > 30 and read depth > 2 using bcftools annotate (82) and VCFfilter (83). Further filtering was performed in PLINK 1.9 (84–86) after which data included only SNVs with minor allele frequency >= 2.5% and 80% genotyping rate. Principal components analysis (PCA) based on the variance-standardised relationship matrix was performed in PLINK 1.9 (84–86) and generated eigenvectors and eigenvalues were visualised in RStudio v2021.9.0.351 (87) using the tidyverse package (88). Dendrograms visualising clustering results from identity-by-state matrix and dissimilarity matrices were created in RStudio v2021.9.0.351 (87). Dendrogram visualising hierarchical clustering performed in RStudio was visualised using FigTree v1.4.4 (89).

### Density and position of single nucleotide variants

The density of SNVs per 100,000 bp, filtered the same way as above, was visualised with CircosVCF (90) with each circle representing an individual plant. The mean density of SNVs per 100,000 bp of the length of each chromosome of the *C. sativa* reference genome (93) was visualised using the RStudio v2021.9.0.351 (87) and tidyverse package (88). Chi-square statistics for the number of SNVs per chromosome between samples were calculated in RStudio v2021.9.0.351 (87) using the tidyverse package (88). Overlaps in positions of SNVs between individual samples were calculated using the VCF Toolz (91).

## Acknowledgements

NT is funded by the UCD School of Biology and Environmental Science. SS was supported by a postdoctoral scholarship of the Irish Research Council and GreenLight Medicines (Ireland) under the Enterprise Partnership Scheme (grant no. EPSPD/2019/220, Enterprise Partner GreenLight Medicines) during part of the project. We are grateful to Jack Forsythe who helped with DNA extraction. We thank Prof. Mali Salmon-Divon from Ariel University who was instrumental for troubleshooting CircosVCF. Open access funding was provided by IReL.

## Author contributions

RM, PFM and SS conceptualised the project. SS performed the experiments. NT and GP performed the data analysis. NT, SS and RM drafted the manuscript. All authors edited and finalised the manuscript.

## Supporting information captions

**S1 Fig. Imaging of hemp plants used for this study.** All hemp plants used for this study were imaged on day 130 after sowing. Individual plants are grouped by cultivar.

**S2 Fig. Hemp plant architectural traits vary within and between cultivars.** The ten hemp cultivars showed differences in morphology and agriculturally important traits. Variability was observed for the number of nodes measured or calculated per 1-metre length (a) and the area of the stem cut ∼5 cm above the soil line at the end of the trial (b). The number of nodes indicates the number of side branches. Significance levels are indicated with letters (p < 0.05).

**S3 Fig. Flower density varies within and between cultivars.** Flower density of female and monoecious hemp plants was recorded at the end of the cultivation period. The size of points increases with the number of individuals with a given flower density.

**S4 Fig. Sex expression of monoecious hemp is not uniform.** Flower sex expression of monoecious hemp cultivars at the end of the cultivation period is plotted as a stacked bar graph, indicating individuals with male and female flowers (dotted), female only flowers (crosshatched) and non-flowering individuals (blank).

**S5 Fig. SNV density plotted for individual cultivars.** The distribution of SNVs plotted on a circos plot (CircosVCF, 79) using the *C. sativa* reference genome (93): ‘Fedora 17’ (a), ‘Felina 32’ (b), ‘Finola’ (c), ‘Futura’ (d), ‘Futura 75’ (e), ‘Georgien’ (f), ‘Kompolti CAN 56’ (g), ‘Kompolti CAN 70’ (h), ‘Korea’ (i), ‘Santhica 27’ (j). Density of filtered SNVs is represented per 100,000 bp.

**S6 Fig. Heterozygosity found in *C. sativa* and other crops**. The heterozygosity of *C. sativa sativa* (this study, 21, 22) in comparison to an accession of a diverse panel of different crops including cultivars and landraces (Plant Imputation database, (75)). The comparability of hemp data is limited because the amount of individuals of other crops was nearly thousandfold higher.

**S7 Fig. Principal component analysis based on SNVs for factors PC2 and PC3**. Principal components from the PCA show clustering of individuals that belong to the same hemp cultivar. PC1 and PC3 (a), PC2 and PC3 (b). For PC1 vs PC2 see Figure 5a.

**S8 Fig. Hierarchical clustering of individual hemp plants based on SNVs**. Hierarchical cluster dendrogram based on pairwise identity-by-state (IBS) values (a) and a dendrogram based on hierarchical hclust clustering values (b) from SNV data of all sequenced individuals show a clustering of individuals that belong to the same hemp cultivar.

**S1 Table**. Detailed phenotypic and developmental traits of ten hemp cultivars grown under greenhouse conditions.

**S2 Table**. The number of SNVs per individual hemp plants.

**S3 Table**. Heterozygosity of hemp cultivars calculated as a ratio of heterozygous to homozygous alternative alleles for hemp cultivars published here, in Sawler *et al.* (2015) (21) and Lynch *et al.* (2016) (22).

